# In vivo mapping of tissue- and subcellular-specific proteomes in *Caenorhabditis elegans*

**DOI:** 10.1101/066134

**Authors:** Aaron W. Reinke, Raymond Mak, Emily R. Troemel, Eric J. Bennett

## Abstract

Multicellular organisms are composed of tissues that have distinct functions requiring specialized proteomes. To define the proteome of a live animal with tissue and subcellular resolution, we adapted a localized proteomics technology for use in the multicellular model organism *Caenorhabditis elegans*. This approach couples tissue- and location-specific expression of the enzyme ascorbate peroxidase (APX), which facilitates proximity-based protein labeling in vivo, and quantitative proteomics to identify tissue- and subcellular-restricted proteomes. We identified and localized over 3000 proteins from strains of *C. elegans* expressing APX in either the nucleus or cytoplasm of the intestine, epidermis, body wall muscle, or pharyngeal muscle. We also identified several hundred proteins that were specifically localized to one of the four tissues analyzed or specifically localized to the cytoplasm or the nucleus. This approach resulted in the identification of both previously characterized and unknown nuclear and cytoplasmic proteins. Further, we confirmed the tissue- and subcellular-specific localization of a subset of identified proteins using GFP-tagging and fluorescence microscopy, validating our in vivo proximity-based proteomics technique. Together, these results demonstrate a new approach that enables the tissue- and subcellular-specific identification and quantification of proteins within a live animal.

## Introduction

Animal development and function relies on the coordinated expression of proteins in specific tissues and the correct localization of those proteins to specific subcellular compartments. Understanding both the tissue and the subcellular localization of a protein can be critical to revealing its function. Because of the fundamental importance of understanding protein localization, several experimental approaches have been established to globally define tissue- and compartment-specific protein expression. One approach relies on generating fluorescent protein fusions with proteins of interest and analyzing protein localization using microscopy. A seminal study using the single-celled yeast *Saccharomyces cerevisiae* determined the subcellular localization of a majority of the proteome^1^. This fluorescent tagging approach has been subsequently applied at the genome scale to the bacteria *Escherichia coli* and *Caulobacter crescentus*^2–4^. Due to the cellular complexity in animals, global determination of protein localization has been more challenging. Genome-wide fluorescent tagging approaches have been initiated to analyze protein localization in the animals *Caenorhabditis elegans* and *Drosophila melanogaster*. These efforts are impressive, but due to the large number of proteins and difficulty in generating transgenic animals, the most comprehensive attempts so far have only localized 12 percent of the interrogated proteome.^5,6^.

Another approach to define tissue and subcellular-specific proteomes relies on biochemical isolation of tissues followed by mass-spectrometry to identify proteins. This approach has been widely utilized to define large-scale tissue maps for the human and mouse proteomes based on dissection of specific organs followed by mass spectrometry analysis^7,8^, but these studies lack cellular resolution within tissues, which can be comprised of multiple different cell types^9^. In addition, these studies are difficult to perform with small organisms where tissue dissection is challenging, such as with *C. elegans*. Cellular compartment-specific proteomes have also been generated, but biochemical isolation techniques can result in a loss of integrity of isolated subcellular compartments leading to incomplete or inaccurate spatial information^10^.

To address the drawbacks associated with tissue dissection, a class of approaches have been developed to define protein expression in the individual tissues and cells of live animals. These approaches rely on the tissue-specific expression of a modified tRNA synthetase that selectively incorporates unnatural amino acids with chemical handles into the proteome. The labeled proteins can then be purified and mass spectrometry used to identify proteins from specific tissues. Several studies have successfully used this approach in *C. elegans* and *D. melanogaster*^11,12^. Although this represents a promising class of approaches, these types of methods are limited to all proteins expressed in a tissue and do not allow for direct detection of protein subcellular localization. Additionally, these approaches can have limited sensitivity since the incorporation of the unnatural amino acid is reported to be less than 1 percent per codon^12^.

Recently, a proteomic technique that labels proteins in discrete locations of live cells without the need for biochemical fractionation has been developed. Known as spatially restricted enzymatic tagging, this method allows for proteins in specific cellular compartments to be tagged with a chemical handle in vivo^13^. This approach relies on the localized expression of soybean ascorbate peroxidase (APX), which in the presence of hydrogen peroxide (H2O2) and biotin-phenol, catalyzes the formation of a phenolic radical that can covalently modify proximal proteins with biotin. These proteins can then be isolated with streptavidin beads and then identified and quantified using mass spectrometry. The efficacy of this method was demonstrated in human cells, where APX was shown to be active in a large number of subcellular compartments^13^. This approach was also recently utilized to identify proteins localized to the mitochondria in dissected fly tissues^10^. These studies demonstrate the potential of this technique to be used to identify proteins from live animals.

The nematode *C. elegans* offers an ideal system for application of spatially restricted enzymatic tagging in a live animal. *C. elegans* is a simple multicellular model organism that has only 959 cells organized into conserved tissues such as muscle and intestine^14^. *C. elegans* has provided the basis for fundamental discoveries in signaling, development and neurobiology, but is lacking a global description of protein localization in its various tissues^15^. To generate a tissue and subcellular-specific map of protein localization in *C. elegans*, we expressed APX localized to two subcellular compartments (cytoplasm and nucleus) in each of four tissues (intestine, epidermis, body wall muscle, and pharyngeal muscle). Subsequent isolation of biotinylated proteins and identification by quantitative mass spectrometry allowed us to quantitatively compare proteins detected in the cytoplasm and nucleus within each tissue to provide a catalog of protein expression specific to either subcellular compartment within specific tissues. Together, these results demonstrate a global approach to characterize tissue- and compartment-specific proteomes in vivo.

## Results

### Development of spatially restricted enzymatic tagging in the intestine of *C. elegans*

To develop a system for identifying proteins with a high degree of temporal, tissue, and subcellular resolution in a live animal, we adapted the use of spatially restricted enzymatic tagging to the nematode *C. elegans*, focusing first on the intestine (Fig. 1). We first generated transgenic animals that express ascorbate peroxidase (APX) as a single genomic copy in *C. elegans* using the MosSCI method^16^. The enzyme was fused to GFP for visualization, localized to the cytoplasm using a nuclear export signal (NES), and specifically expressed in the intestine using the *spp-5* promoter. We confirmed the intestinal cytoplasmic-specific expression of this strain, GFP-APX-NES, by fluorescent microscopy (Fig. S1). As a negative control we also generated a strain in a similar manner that expresses GFP in the intestine without the APX enzyme.

**Figure 1.**
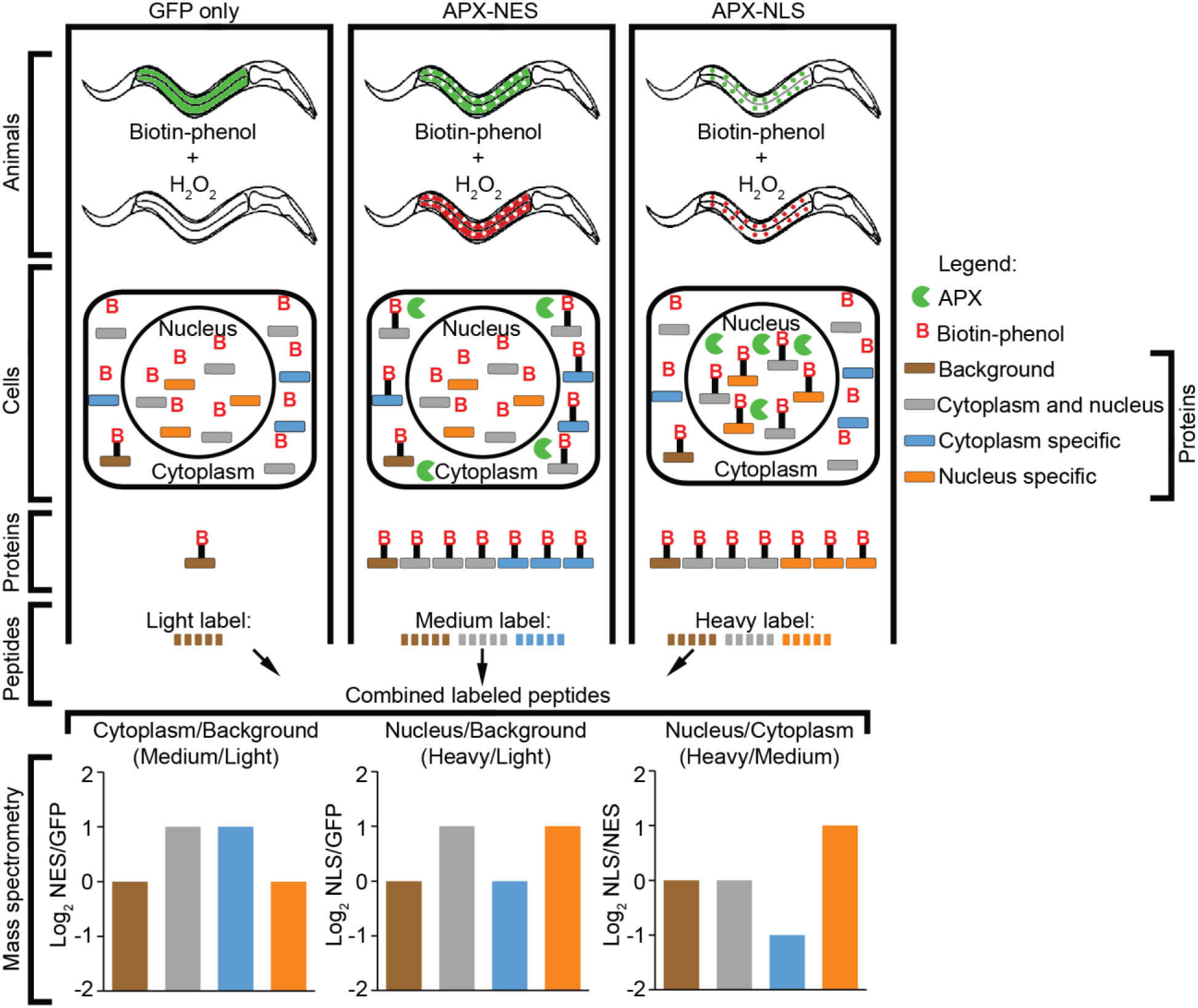
Overview of approach to identify tissue and subcellular-specific protein expression in *C. elegans*. Schematic of spatially restricted enzymatic tagging in *C. elegans*. Animal strains are generated that express the APX enzyme in either the cytoplasm or the nucleus in a tissue-specific manner, such as in the intestine as illustrated here. Animals are treated with biotin-phenol that diffuses into cells. The APX enzyme, in presence of H_2_O_2_ and biotin-phenol catalyzes the formation of a phenoxyl radical that covalently labels neighboring proteins with biotin (red B with black bars)^13^. Thus, biotin labeling (labeled in red) of proteins occurs in whichever specific tissues and subcellular locations the enzyme is expressed. Three strains are used to measure protein localization in each tissue: GFP only, GFP-APX-NES, and GFP-APX-NLS. Spatially restricted tagging is performed using these three strains and then the proteins are extracted and purified using streptavidin beads. These purified proteins are then digested into peptides and labeled using reductive dimethyl labeling for quantitative comparisons between the three strains. The peptides from each sample are then combined and peptide ratios in each sample are measured using mass spectrometry. The peptide ratios can then be used to determine if the protein is detected over background and if it is enriched in either the nucleus or cytoplasm.

To test the activity of APX in *C. elegans*, we grew a single plate of 30,000 synchronized animals on bacteria until the fourth larval (L4) stage (44 hours) at 20 °C. Populations of animals were grown for both the strain expressing APX and the negative control strain expressing GFP without APX. These animals were then washed off the plate and treated with the biotin-phenol substrate for 1 hour, followed by the addition of H2O2 for 2 minutes. The labeling reaction was then quenched and proteins from these animals were extracted and purified with streptavidin beads. Proteins were then eluted from the beads, separated on SDS-PAGE gels, and visualized with Oriole staining. In the absence of APX expression, endogenously biotinylated proteins were detected^17^. The presence of APX did not result in the expected increase of biotinylated proteins compared to the control GFP only expressing animals (Fig. 2A).

**Figure 2.**
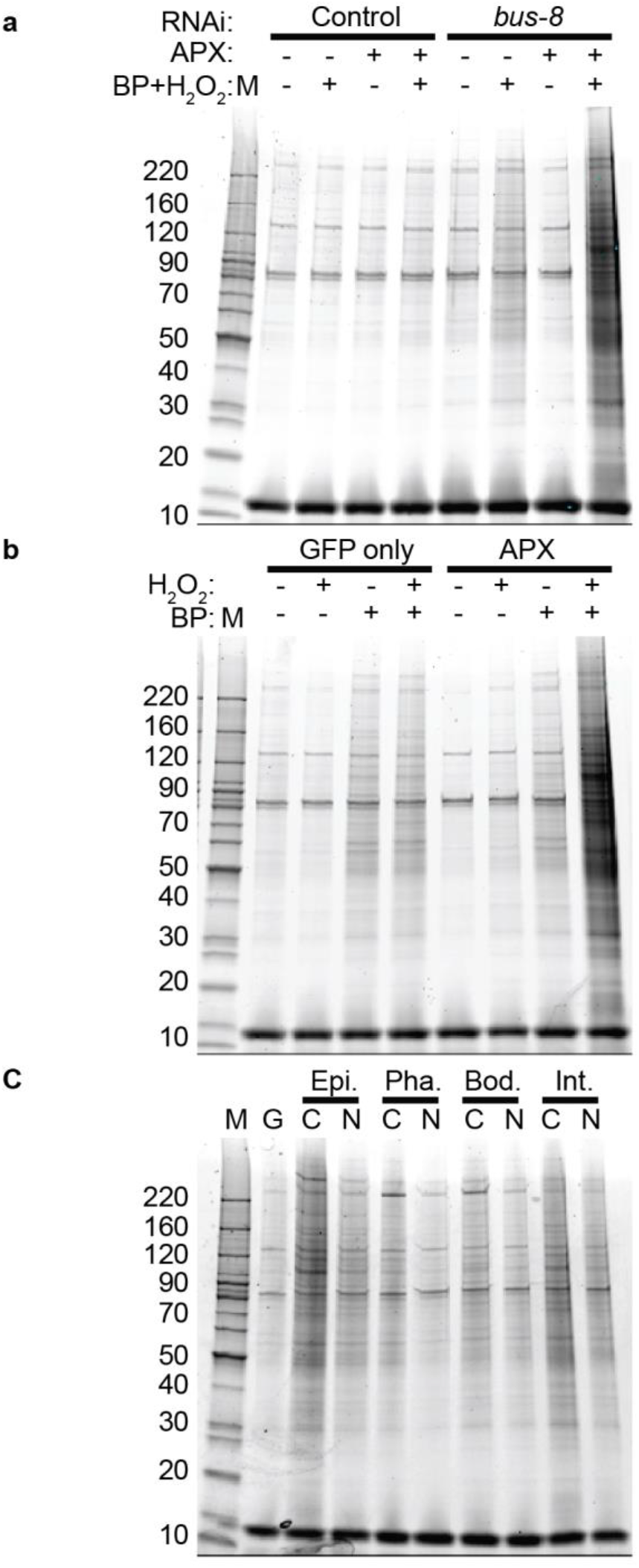
Efficient spatially restricted enzymatic tagging in *C. elegans* is dependent on biotin-phenol, H2O2, and *bus-8* RNAi. **a-c**. Streptavidin-purified proteins from *C. elegans* protein extracts were visualized with the Oriole stain. **a**. Animals expressing GFP-APX (APX +) in the intestinal cytoplasm or a GFP only control strain (APX −) were grown on plates with either control (L4440) or *bus-8* RNAi. Animals were either treated or untreated with biotin-phenol (BP) and H2O2. Protein markers are indicated (labeled M). **b**. Animals expressing GFP-APX in the intestinal cytoplasm or a GFP control strain grown on plates with *bus-8* RNAi were treated with either H2O2 or biotin-phenol, or both. c. Strains of *C. elegans* expressing GFP-APX in either the cytoplasm (C) or nucleus (N) of the epidermis (Epi.), pharyngeal muscle (Pha.), body wall muscle (Bod.), or intestine (Int.). A strain expressing GFP only (G) is the negative control. All strains were grown on plates with *bus-8* RNAi and treated with biotin-phenol and H_2_O_2_.

Based on a lack of APX-mediated biotin labeling and our observation that APX was properly expressed, we hypothesized that the concentration of the biotin-phenol substrate in the worm intestinal cells was inadequate to facilitate efficient protein labeling. To investigate whether labeling efficiency could be improved by increasing the permeability of the *C. elegans* cuticle to biotin-phenol, we knocked down the expression of the *bus-8* gene. BUS-8 is a glycosyltransferase involved in cuticle development, and reduction of BUS-8 function has been shown to increase small molecule permeability by decreasing cuticle integrity^18^. Therefore, we fed animals bacteria that express double-stranded RNA against *bus-8* to induce RNAi knockdown of *bus-8* expression. Under these conditions, APX-expressing animals treated with the biotin-phenol substrate displayed increased biotin tagging of proteins compared to control animals that do not express APX (Fig. 2A). Therefore, decreasing cuticle integrity appears to increase availability of the substrate, leading to increased biotinylation of cellular proteins by APX. Because of this substantial increase in biotinylation efficiency, we performed all subsequent experiments by growing animals on bacteria expressing the *bus-8* RNAi clone to increase cuticle permeability and APX-mediated protein biotinylation.

We then investigated whether the biotinylation reaction described above depends upon the previously described components for an APX-mediated reaction: the APX enzyme, the biotin-phenol, and H2O2. First, we observed a slight increase in background biotinylation when the biotin-phenol was added to the control animals that do not express APX (Fig. 2B), indicating that a low level of biotinylation occurs without the enzyme. However, there was a substantial increase in labeling when animals expressing APX were exposed to biotin-phenol and H2O2 (Fig. 2B), demonstrating that efficient biotinylation is greatly potentiated by APX expression. We also found that both biotin-phenol and H2O2 were required for efficient labeling (Fig. 2B), consistent with APX-mediated biotinylation in humans cells also being dependent on H2O2. Notably, in our experiments animals are incubated with biotin-phenol for 1 hour, while H2O2 is only added for a period of 2 minutes before being quenched. Because H2O2 is required for labeling, this result indicates that the labeling reaction is rapid and occurs within only 2 minutes in *C. elegans*.

### Biotin labeling of proteins in specific locations within *C. elegans*

To determine whether protein biotinylation and analysis could be performed in other tissues and compartments in *C. elegans*, we created a panel of strains expressing the APX enzyme in different tissues and subcellular locations. In addition to a version of the protein localized to the cytoplasm, we created another version where APX is localized to the nucleus using a nuclear localization signal (NLS). In addition to the intestine, we expressed the enzyme in three other tissues using the following tissue-specific promoters: epidermis (*dpy-7*), body wall muscle (*myo-3*), and pharyngeal muscle (*myo-2*) (Table S1). In total, we generated eight strains expressing the APX enzyme. This panel of strains and the negative control strain were grown and treated with biotin-phenol and H2O2 to label proteins as described above. The APX-mediated biotinylation of proteins in each of these tissues was examined, revealing a clear increase in biotinylation in every location compared to the negative control strain, with the exception of the nuclear localized enzyme in the pharyngeal muscle (Fig. 2C). Of the tissues analyzed in our panel, this location represents the smallest tissue and thus it is likely that labeled proteins could not be detected over background biotinylation levels.

To confirm the location specificity of the labeling, we used fluorescence microscopy to visualize where within the animal proteins were biotinylated. After treatment of the animals with biotin-phenol and H2O2, we fixed and stained the animals with anti-GFP antibodies to localize the fusion protein and with fluorescent streptavidin to localize biotinylated proteins. Fluorescent microscopy was used to analyze these stained animals and we found that the location of the biotin labeling was dependent on the tissue and compartment where APX was expressed (Fig. 3). Although labeling could not be detected on gels when the enzyme was localized to the nucleus of the pharyngeal muscle, efficient and specific labeling was observed in this location by microscopy (Fig. 3). The control GFP only strain lacking APX did not display biotin labeling, demonstrating the specificity of the approach (Fig. 3). The combined results from examining protein extracts and from microscopy indicate the ability to label proteins in each of the eight locations that we tested. Together this demonstrates the efficacy of in vivo spatially restricted enzymatic tagging in *C. elegans*.

**Figure 3.**
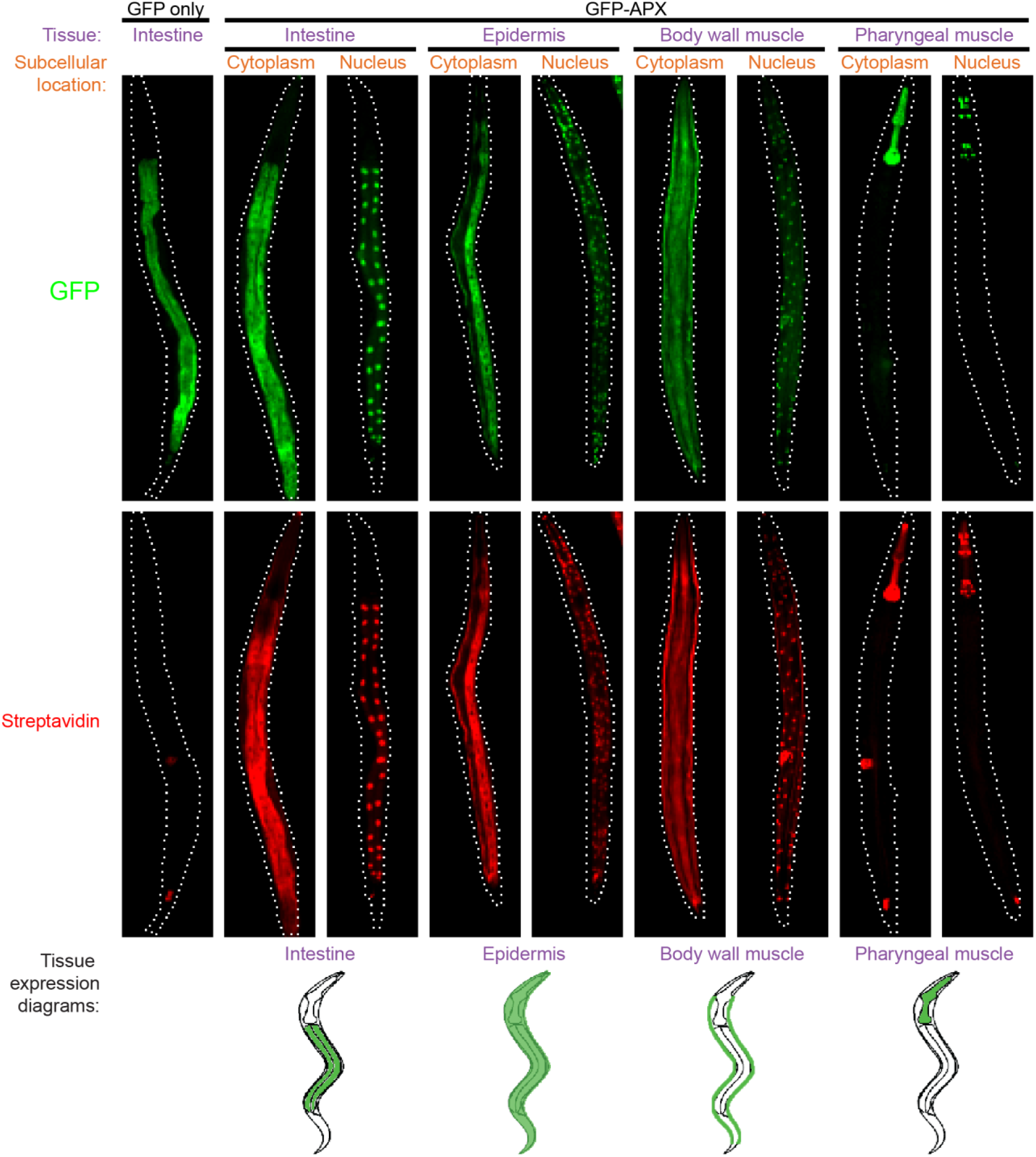
APX-mediated biotin labeling in vivo displays tissue and subcellular specificity. Spatially restricted enzymatic tagging was performed on strains of *C. elegans* expressing APX in the cytoplasm or nucleus of the intestine, epidermis, body wall muscle, or pharyngeal muscle as indicated. A strain expressing GFP without APX was used as a negative control. Animals were fixed and stained for GFP (top, green) to determine the localization of the enzyme, and streptavidin (middle, red) to determine the location of protein biotinylation. Representative images are shown for each strain. Animals are aligned so that the anterior is up. Tissue expression diagrams show the location of each tissue in *C. elegans* (Bottom).

### Identification of *C. elegans* cytoplasmic and nuclear proteins expressed in specific tissues using mass spectrometry

Having confirmed the tissue and cell compartment specificity of our in vivo APX-mediated proximity tagging approach, we then set out to identify proteins that are present in the cytoplasm and nucleus and in specific tissues using mass spectrometry. We employed a quantitative strategy to rapidly and accurately compare proteins isolated from each tissue and between different subcellular locations. To identify cytoplasmic and nuclear proteins, we used a set of three strains for each tissue. We utilized the strains expressing APX-NES and APX-NLS in each tissue and the negative control strain that expresses GFP without APX in the intestine. (Fig. 1). For the three samples in each tissue set, proteins were labeled by APX-mediated biotinylation and isolated as described above. Proteins bound to streptavidin-agarose from the three strains (GFP-APX-NES, GFP-APX-NLS, and GFP only) were then digested with trypsin.

For quantitative comparisons among the three samples in each tissue set, peptides from each sample were labeled with a different isotopic tag using reductive dimethyl labeling (Fig. 1)^19^. For each tissue set, differentially-labeled reductive dimethylated peptides from each sample were mixed in equal proportion prior to analysis by high-resolution mass spectrometry. Samples from each tissue set were prepared and analyzed in triplicate. To evaluate the ability of this approach to identify proteins above background levels, we initially performed control experiments using the three strains of the intestine tissue set (GFP-APX-NES, GFP-APX-NLS, and GFP only). As a control, we pooled peptide samples from each of the three strains, separated them into three identical pools prior to dimethyl labeling, and upon remixing in equal amounts found that less than 5% of all proteins displayed quantitative ratios greater than 2-fold between pools establishing a base false discovery rate for our experimental method (Fig. S2A). In contrast, when peptide samples from the three strains were labeled by reductive dimethyl labeling individually before mixing, over 90% of proteins from the APX-NES and APX-NLS samples had ratios over the GFP only control sample greater than 2-fold (Fig. S2B). These control experiments demonstrate the effectiveness of using reductive dimethyl labeling and mass spectrometry to quantitatively identify APX-mediated biotinylated proteins and established a 2-fold threshold as being able to differentiate proteins above background which we employed in subsequent analyses.

To identify proteins that are present in the eight locations we analyzed, ratios of proteins identified from the APX-NES and APX-NLS strains for each tissue were compared to the GFP-only control strain. We evaluated the reproducibility of our approach by comparing how many proteins with ratios greater than 2-fold over background were identified in at least one replicate compared to those identified in all three replicates (Fig. S3). In the largest locations measured, the cytoplasm of the intestine and epidermis, over 60% of proteins that were identified 2-fold above background in one replicate were also identified as 2-fold above background in the two other replicates. In contrast, in the smallest location measured, the pharyngeal nucleus, only 3% of proteins identified 2-fold above background in one replicate were also 2-fold above background in the two other replicates. These differences are largely due to the variability in the number of proteins identified between replicates. For example, the number of proteins identified in the intestinal cytoplasm were between 2376 to 2575, while for the pharyngeal nucleus the range was much larger, with between 116 to 759 proteins identified (Fig. S3).

We then identified proteins from each of the experimental strains using the criteria that each protein has a ratio at least 2-fold over background in 2 of the 3 biological replicate experiments (see methods). Using these criteria, we identified between 108 to 2484 proteins for each of the 8 strains (Fig. 4A). The proteins identified and ratios between pairs of strains in the same tissue set are reported in Table S2. The largest tissues, the intestine and epidermis, had the most identified proteins, followed by the body wall muscle and then the pharyngeal muscle. More proteins were identified from the cytoplasm than from the nucleus for each tissue. A total of 3187 proteins were identified in at least one of the eight strains (Fig 4B).

**Figure 4.**
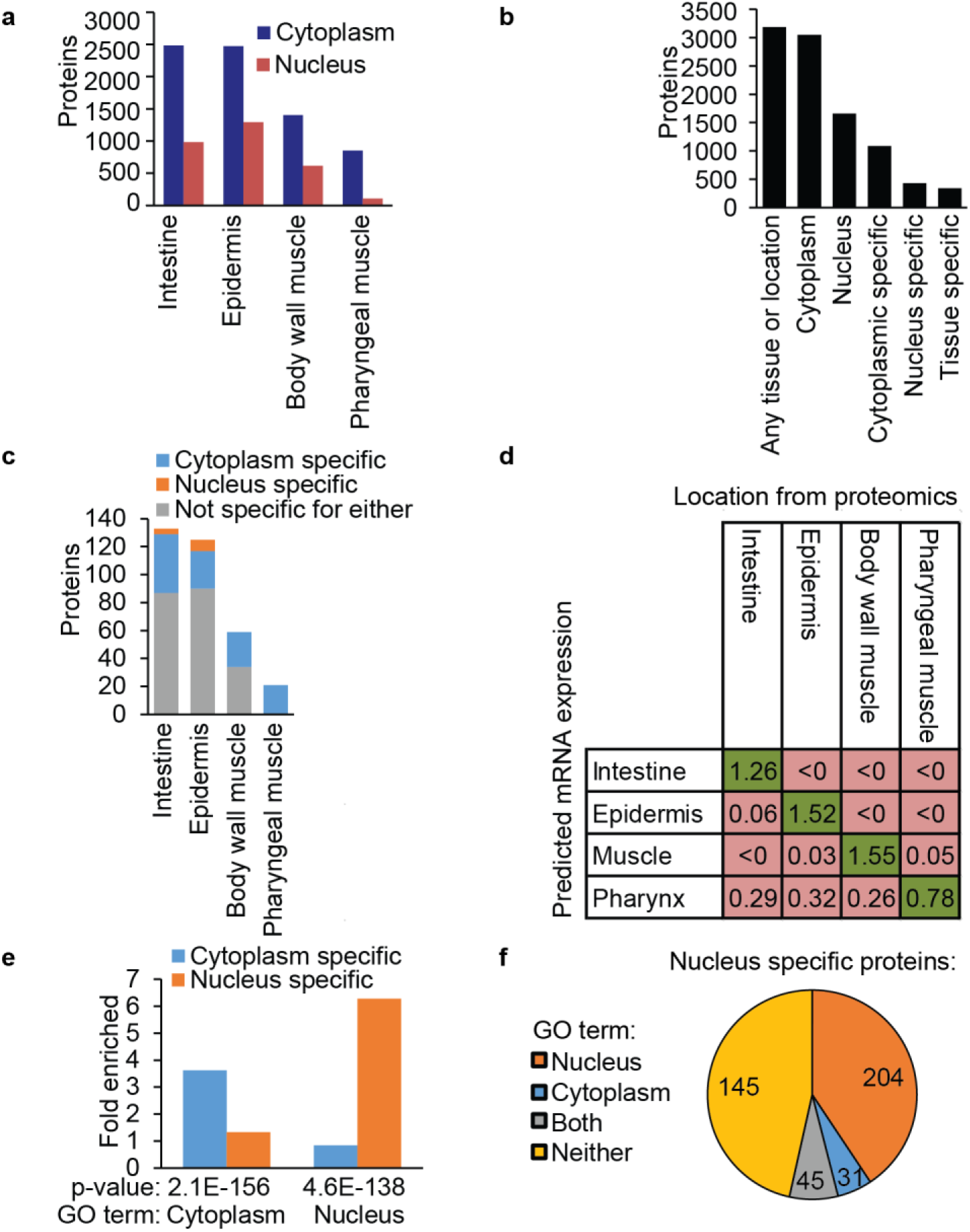
Identification of *C. elegans* proteins with tissue- and subcellular-specific localizations. **a**. The number of proteins identified above background from the tissue or subcellular location for each of the eight locations indicated. **b**. The total number of proteins identified in our experiments that were detected in different locations or that were detected as being specific to a location. **c**. The number of proteins we identified in the indicated tissue, but not in the other three tissues. For each tissue, three categories of proteins are shown, those that are specific to the cytoplasm (orange), those that are specific to the nucleus (blue), and those that are not specific to either compartment (grey). **d**. Comparison of the identified tissue-specific proteins to a dataset of predicted mRNA expression ^20^. The data presented are the average of all the mRNA expression prediction scores for each tissue-specific protein in each of the four tissues. Higher prediction scores are more likely to be expressed in that tissue. Each column represents proteins identified as specific to that tissue, compared to the predicted mRNA expression of the tissue in each row. The highest average score in each column is shaded in green, and all other scores in the column are shaded in red. **e**. GO term enrichment analysis of proteins identified in our experiments as specific to either the cytoplasm or the nucleus. **f**. Pie chart of the nucleus-specific proteins we identified in our experiment, and whether they have a GO term location of either the nucleus, cytoplasm, both, or neither.

### Identification of tissue and subcellular specific *C. elegans* protein expression

We then investigated which of the proteins that we identified displayed tissue-specific expression. To identify proteins that are tissue-specific, we compared proteins that were detected as being 2-fold above background in one of the four tissues in two of the three replicates and not detected 2-fold above background in any of the replicates in the other three tissues. This resulted in the identification of 338 proteins that were specific to only one tissue (Fig. 4C). To assess the accuracy of these proteins identified as tissue-specific, we compared them to an existing comprehensive data set of tissue-specific mRNA expression^20^. This data set was generated from experimental expression data and prediction scores were made for each gene in each of the four tissues we tested. The set of proteins we identified as being specific for each tissue had the highest average mRNA expression scores in that tissue compared to the other three supporting the accuracy of our technique, tissues (Fig. 4D). We identified the largest number of proteins from the intestine and epidermis as being tissue-specific. These tissues also had the lowest mRNA expression scores for the proteins identified as being specific to other tissues. We identified the fewest proteins from the pharyngeal muscle as being tissue-specific and this tissue also had higher mRNA expression scores for the proteins identified as being specific to other tissues. This lowered accuracy is likely due to identifying fewer total proteins in the pharyngeal muscle compared to the other tissues (Fig. 4A). Thus some proteins that we describe as specific to non-pharyngeal tissues may actually be expressed in the pharyngeal muscle as well, but we failed to detect them in the pharyngeal muscle due to the lowered sensitivity for our technique in this small tissue. This result suggests these analyses are more useful for the large tissues where we identified more proteins. However, because the pharyngeal-specific proteins we describe do have the highest expression score in the pharynx in comparison to the other tissues, it demonstrates that this analysis is still useful for identifying tissue-specific proteins even in smaller tissues.

We then examined the subcellular specificity of the proteins we identified by comparing the ratios of protein levels in the nucleus compared to protein levels in the cytoplasm. We used the quantitative ratios between the APX-NES and APX-NLS samples to find proteins that were 2-fold enriched in either sample (Fig. S4). This comparison resulted in the identification of 1087 proteins specific to the cytoplasm and 428 proteins specific to the nucleus (Fig. 4E). To assess the accuracy of these location assignments we performed GO term enrichment analysis using PANTHER^21^. Here we found that proteins we identified as being cytoplasm-or nucleus-specific were highly enriched for proteins previously annotated to be cytoplasmic or nuclear (Fig. 4E). Of the 428 proteins identified as nucleus-specific, 176 were not previously known to be nuclear localized (Fig. 4F). We also examined the compartment specificity of the proteins we classified as being tissue-specific and identified 115 of these proteins as being cytoplasm-specific and 12 as being nucleus-specific (Fig. 4C).

To validate our results, we used fluorescence microscopy to confirm the localization of several proteins that we identified to be expressed in specific tissues or subcellular locations using our quantitative proteomics approach. We chose 7 proteins that we measured with high confidence to be either nucleus-specific, cytoplasm-specific, or specific to one of the four tissues (Fig. S5). These seven proteins had no previous experimentally determined location described in WormBase (http://www.wormbase.org). Using TransgeneOme^5^ constructs that contain C-terminal GFP fusions of each protein expressed under the native promoter, we generated strains of transgenic animals overexpressing each protein. We found that each of these seven test proteins localized to the corresponding tissue or subcellular location that we identified by spatially restricted enzymatic tagging (Fig. 5). These results confirm the accuracy of our approach, and demonstrate the efficacy of using in vivo proximity-based labeling methods and quantitative mass spectrometry to identify proteins with tissue-specific and/or subcellular compartment-specific localization. Overall, we present a robust method that can be applied to detect in vivo protein localization in an unbiased manner within intact animals and provide a resource of proteins with specific locations in *C. elegans* (Table S2).

**Figure 5.**
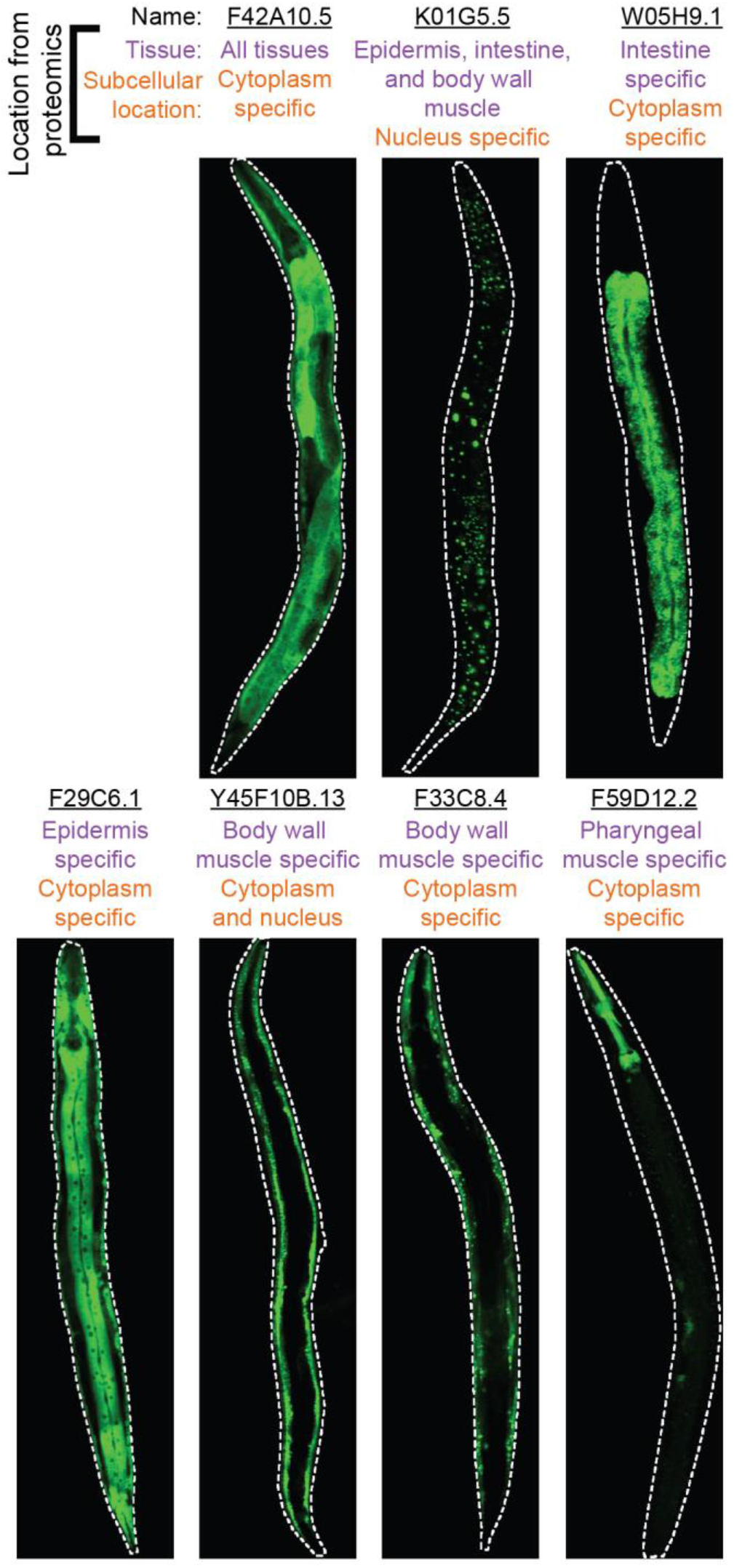
Validation of identified protein locations using fluorescently-tagged proteins. Strains of transgenic *C. elegans* expressing GFP-tagged proteins identified to be tissue-specific or location-specific in our study. Animals were grown to the L4 stage and representative images displaying protein localization are shown. The protein name is listed above each construct-expressing strain. The tissue and subcellular localization determined from our proteomic data is listed below the protein name. Animals are aligned so that the anterior is up.

## Discussion

Here, we describe an approach that allows for the determination of in vivo protein localization in an intact animal through the use of spatially restriced enzamatic tagging. To our knowledge this is the first in vivo localization of a large number of proteins with subcellular resolution in a live animal. Using spatially restricted enzymatic tagging in *C. elegans*, we provide one of the largest systematic identifications of proteins that have tissue or compartment-specific localizations. We identified a total of 3187 proteins, of which 1673 are localized to a specific tissue or subcellular compartment. This resource represents an advancement from previous studies aimed at identifying protein localization using GFP-tagging, in which the location of 230 proteins were characterized. A different approach using unnatural amino acid labeling in the pharyngeal muscle in *C. elegans* identified 43 proteins that were greater than 2-fold enriched above background levels^11^. In contrast our technique identified 887 proteins total in either the cytoplasm or nucleus of the pharyngeal muscle.

Despite our success in using spatially restricted enzymatic tagging to identify tissue and subcellular specific protein expression, there are some limitations to this approach that warrant discussion. The number of proteins identifed even in the largest tissues is only ~25% of the number of mRNAs demonstrated to be expressed in the same tissue^22^. Additionally, fewer proteins were identified in the smaller tissues than larger tissues, and in the nucleus compared to the cytoplasm. These concerns may be addressed in the future by using recently developed versions of APX that have increased sensitivity^23^. Additionally, as the variability of identified proteins in the smaller tissues was greater (Fig. S3), the measurement of additional replicates could be used to increase sensitivity of protein detection. Another issue is that our approach relies on feeding animals *bus-8* RNAi, which causes improper development of the cuticle and locomotion defects^18^. However, we were able to confirm the specific location of a number of proteins using transgenic analysis in wild-type animals, demonstrating that protein localization in bus-8-defective animals appears to be largely similar to wild-type animals. Additonally, there are potentially other mutants or chemical approaches that could be employed to improve the accessibility of the biotin-phenol substrate and may lessen their physiological impact on the animals.

The methodology we describe here can be expanded upon to obtain greater tissue and subcellular resolution. Spatially restricted enzymatic tagging has also been reported to work in other subcellular locations in human cells including the endoplasmic reticulum, mitochondria, and plasma membrane^13^. We were able to identify proteins from all eight of the locations to which we localized the enzyme. Thus, it is likely that this approach could be applied to a number of other tissues and subcellular locations in *C. elegans*.

This technique we describe also could be useful for a number of applications. This technique could be used to determine protein translocation between the cytoplasm and the nucleus in different growth conditions. Through the use of a yeast GFP-tagged library, 71 cytoplasmic proteins were shown to localize to the nucleus under starvation conditions, demonstrating that large numbers of proteins translocate in response to stress^24^. Using our described approach, these types of translocation events could be measured globally in live animal using a smaller number of strains, to investigate the response to various stress conditions. Although our experiments were focused on quantifying differences between the cytoplasm and nucleus, the quantitative labeling scheme we employed is flexible and could be used to directly compare levels of proteins between different tissues. Additionally, this approach could also be adapted to measure posttranslational modifications, such as phosphorylation. This would allow comparing differences in post-translational modifications of proteins between different compartments and between different tissues^25^. This approach can also be applied to identify proteins from pathogenic or symbiotic microbes that localize to different host tissues and subcellular locations^26^. Spatially restricted enzymatic tagging has now been reported in *D. melanogaster, C. elegans*, and human cells and thus can likely be used in any organism where transgenic techniques exist and biotin-phenol can be delivered.

## Acknowledgements

We thank Keir Balla, Robert Luallen, and Kirthi Reddy for providing helpful comments on the manuscript. We thank Nick Kosa and Robert Ardecky for providing aid in purifying the biotin-phenol. AWR is a Monsanto Fellow of the Life Sciences Research Foundation.

## Author Contributions

AWR designed, conducted, and analyzed experiments and co-wrote the paper. RM performed mass spectrometry analysis. EJB designed experiments, performed mass spectrometry analysis, and co-wrote the paper. ERT provided mentorship and guidance of the project and co-wrote the paper.

## Methods

### Cloning and generation of strains

The protein sequence of soybean ascorbate peroxidase (APX) with the W41F mutation^13^ was optimized for *C. elegans* expression using DNAworks to design primers^27^. Primers were annealed using a two-step PCR method and double-stranded DNA was cloned into Gateway vector pDONR 221 using BPclonase II (Thermo Fisher). This construct was modified using Gibson cloning^28^ with an N-terminal fusion of GFP. This construct was additionally modified to encode the C-terminal nuclear export signal (NES: LQLPPLERLTLD) and nuclear localization signal (NLS: PKKKRKVDPKKKRKVDPKKKRKV) by encoding these tags into primers, amplifying the plasmid with PCR and ligating the PCR product. Upstream regions of the following *C. elegans* genes were used as promoters: *dpy-7* (epidermis) *spp-5* (intestine) *myo-2* (pharyngeal muscle) *myo-3* (body wall muscle). These promoters were cloned into the 5’ plasmid pDONR P4-P1R using BP clonase II (Thermo Fisher). Multisite Gateway cloning was used to generate targeting constructs using LR clonase II plus (Thermo Fisher) to combine one of the four promoter plasmids, one of the two APX containing plasmids, the 3’ plasmid pDONR P2R-P3 vector containing the 3’ region of *unc-54*, and the destination vector pCFJ150. These targeting constructs along with the Mos1 transposase and marker plasmids were injected into *unc-119* mutants from the strain EG6699^29^. Non-Unc worms were recovered and each transgenic strain was backcrossed into the wild-type N2 strain 3 times. The homozygote was used in subsequent experiments. Transgenic strains expressing transgeneOme GFP-tagged proteins as extrachromosomal arrays were generated by injection of constructs into EG6699 and selecting non-unc animals. The following amount of DNA was injected for each construct: 100 ng/μl of each construct for F33C8.4, Y45F10B.13, F59D12.2, and F29C6.1, 50 ng/μl of the construct with 50 ng/μl pBSK for F42A10.5, and 10 ng/μl of the construct with 90 ng/μl pBSK for W05H9.1 and K01G5.5. All strains used in the study are listed in Table S1. All *C. elegans* strains were maintained using standard procedures^30^.

### Spatially restricted enzymatic tagging in *C. elegans*

Populations of animals were grown and bleached to recover eggs, which were then hatched to generate first larval stage (L1) synchronized animals^30^. ~30,000 L1 animals of each strain in 2.5 ml M9 buffer were added to a 15 cm RNAi plate seeded with HT115 bacteria expressing a *bus-8*RNAi feeding clone^31^. Animals were protected from light and grown to the L4 stage on these plates for 44 hours at 20°C. To recover animals, each plate was washed with M9T (M9/0.1% Tween-20). The recovered animals were washed once with M9T. These animals were then placed into 1.5 ml tubes in a total of 100 μl M9T. T o each sample was added 900 μl of labeling solution (0.1 % Tween-20, M9, and 3.3 mM biotin-phenol, synthesized as previously described)^13^. Samples were incubated for 1 hour at 22-24°C on an end-over-end rotator. To activate biotin labeling, 10 μl of 100 mM H_2_O_2_ was added for 2 minutes. To quench the reaction, 500 μl quench buffer (M9/0.1% TWEEN-20/ 10 mM sodium azide/10 mM sodium ascorbate/ 5mM Trolox) was added. Samples were then washed with 1 ml of quench buffer 4 times. After the last wash, the remaining buffer was removed and 800 μl lysis buffer (150 mM NaCl/50 mM Tris pH 8.0/1% TritonX-100/0.5% sodium deoxycholate/0.1%SDS/10 mM sodium azide/ protease complete tablet (Roche)/10 mM sodium ascorbate/ 5mM Trolox/1 mM PMSF) was added. Animals were then immediately frozen dropwise in liquid N_2_.

To extract proteins, frozen worm pellets were ground to a fine powder in liquid N_2_. To generate supernatants these protein extracts were then centrifuged for 10 min at 21,000 × *g* at 4 °C. The supernatant was then filtered over a desalting column with a 7 kilodalton molecular weight cutoff (Pierce). The protein concentrations of the extracts were measured using a Pierce 660 nm Protein Assay and normalized. To 450-550 μg of each sample was added 25 μl of high capacity streptavidin agarose resin (Pierce) in a total of 700 μl lysis buffer. Extracts were incubated with beads for 1 hour on an end-over end rotator. Beads were then washed 5 times with 1 ml lysis buffer, 3 times with 1 ml 8M urea/10 mM Tris pH 8.0, and 3 times with 1 ml PBS. The liquid was removed from the beads and 100 μl of 0.1 μg/μl trypsin (Promega)/100 mM TEAB was added to each sample and incubated at 37°C for 24 hours.

These peptides were then differentially labeled with reductive dimethyl labeling as previously described^19^. Briefly, each of the three samples in the set was labeled with a different isotopic tag that differed by four daltons. To generate samples with a light tag, 4 μl of 4% (v/v) CH2O and 4 μl 600mM NaBH3CN was added. The other tags were generated in a similar way, with CD2O and NaBH3CN being used for the medium tag and C^13^D2O and NaBD3CN being used for the heavy tag. Samples were then incubated for one hour with mixing on an end-over-end rotator. To quench the reaction, 16 μl of 1% (v/v) ammonia was added to each sample. The samples were then acidified by adding 8μl of formic acid. The three samples in each set were then combined.

### Gel analysis of biotinylated proteins

From samples prepared as described above, 15% of beads were removed before digestion. Liquid was removed from the beads and 20 μl of Laemmli buffer with 2 mM biotin was added. Samples were heated for 10 min at 95°C. 15 μl of each sample was loaded onto a 4-20% polyacrylamide gradient gel (Bio-rad). Gels were then stained with Oriole fluorescent gel stain (Bio-rad) to visualize proteins.

### Microscopy

To analyze biotin labeling with immunohistochemistry, worms were fixed using Bouin’s tube fixation method^32^. Fixed worms were stained overnight with 1:500 anti-GFP mouse antibody (Roche) in block buffer (phosphate buffered saline/0.5% TritonX-100, 1% Bovine serum albumin). Worms were then washed with block buffer and stained overnight with 1:500 FITC-conjugated anti-mouse secondary antibody (Invitrogen) and 1:500 streptavidin Alexa Fluor 568 (Thermo Fisher) in block buffer. Worms were then washed in block buffer and imaged using a Zeiss LSM700 confocal microscope. Live worms expressing GFP-tagged TransgeneOme constructs^5^ were grown to the L4 stage, treated with 1 mM levamisole and then imaged as described above.

### Sample analysis by mass spectrometry

Prior to analysis by LC-MS/MS, peptides were desalted by solid-phase extraction using in-house prepared C18 StageTips^33^ and reconstituted in 5% formic acid and 5% acetonitrile (ACN). All samples were analyzed in triplicate using a Q Exactive mass spectrometer (Thermo Scientific, San Jose, CA). The following is a generalized nHPLC and data acquisition method that is representative of individual analyses. Peptides were first separated by reverse-phase chromatography using a fused silica microcapillary column (75 μm ID, 18 cm) packed with C18 silica (ReproSil-Pur 120 C18-AQ, 1.9 μm, Dr. Maisch GmbH) using an in-line nano-flow EASY-nLC 1000 UHPLC system (Thermo Scientific). Peptides were eluted over a 100 minute 0-30% ACN gradient, followed by a 5 minute 30-60% ACN gradient, a 5 minute 60-95% ACN gradient, with a final 10 minute isocratic step at 0% ACN for a total run time of 120 minutes at a flow rate of 250 nl/min. All gradient mobile phases contained 0.1 % formic acid. MS/MS data were collected in data-dependent mode using a top 10 method with a full MS mass range from 400-1800 m/z, 70,000 resolution, and an AGC target of 3e6. MS2 scans were triggered when an ion intensity threshold of 1e5 was reached with a maximum injection time of 60 ms. Peptides were fragmented using a normalized collision energy setting of 25. A dynamic exclusion time of 20 seconds was used and the peptide match setting was disabled. Singly charged ions, charge states above 6 and unassigned charge states were excluded.

### Peptide and protein identification and quantification

The resultant RAW files were converted into mzXML format using the ReAdW.exe (version 4.3.1) program. The SEQUEST search algorithm (version 28) was used to search MS/MS spectra against a concatenated target-decoy database comprised of forward and reversed sequences from the reviewed UniprotKB/Swiss-Prot FASTA *C. elegans* database combined with the UniprotKB E. coli (K12 strain) database, and with common contaminant proteins appended. Each mzXML file was searched in triplicate with the following parameters: 20 ppm precursor ion tolerance and 0.01 Da fragment ion tolerance; Trypsin (1 1 KR P) was set as the enzyme; up to three missed cleavages were allowed; dynamic modification of 15.99491 Da on methionine (oxidation). For searches with light and medium reductive dimethyl labels, additional dynamic modifications of 4.0224 Da on lysine and peptide N-termini and static modifications of 28.0313 Da on lysine and peptide N-termini were included. For searches with light and heavy reductive dimethyl labels, additional dynamic modifications of 8.04437 Da on lysine and peptide N-termini and static modifications of 28.0313 Da on lysine and peptide N-termini were included. For searches with medium and heavy reductive dimethyl labels, additional dynamic modifications of 4.02193 Da on lysine and peptide N-termini and static modifications of 32.05374 Da on lysine and peptide N-termini were included. Peptide matches were filtered to a peptide false discovery rate of 2% using the linear discriminant analysis. Proteins were further filtered to a false discovery rate of 2% and peptides were assembled into proteins using maximum parsimony and only unique and razor peptides were retained for subsequent analysis. All peptide Heavy:Light, Medium:Light, and Heavy:Medium ratios with a signal to noise ratio above 5 were used for assembled protein quantitative ratios.

### Analysis of mass spectrometry data

We classified a protein as being identified above background if it had a greater than 2-fold ratio of NLS or NES over the GFP only strain in 2/3 replicates for a given tissue. The requirements for a protein being classified as being tissue-specific were 1) having a greater than 2-fold ratio of NLS/GFP in 2/3 replicates or a greater than 2-fold ratio of NES/GFP in 2/3 replicates in one of the tissues and 2) not detected in any of the other tissues with greater than a 2-fold ratio of NLS/GFP or NES/GFP in any of the three replicates. These identified proteins were compared to predicted mRNA expression scores^20^. Cytoplasm-specific or nucleus-specific proteins were determined by comparing the ratio between the NES and NLS samples. Because more peptides were detected in the NES sample than the NLS sample from each tissue, the ratios were normalized by the number of peptides identified in each sample. Those with a greater than 2-fold NLS/NES adjusted ratio were classified as nucleus-specific. Those with a greater than 2-fold NES/NLS adjusted ratio were classified as cytoplasm-specific. GO term enrichment was performed using PANTHER^21^. Proteins to validate tissue-specific expression using GFP-tagged constructs were selected based on the criteria of 1) having a greater than 2-fold ratio of NLS/GFP in all three replicates or a greater than 2-fold ratio of NES/GFP in all three replicates and 2) not having NLS/GFP or NES/GFP ratios greater than 0 in any of the replicates for the other three tissues. The proteins to validate as being subcellular specific were chosen based on having NES/GFP and NES/NLS ratios greater than 2-fold in all three replicates in at least three tissues for being cytoplasm-specific and having NLS/GFP and NLS/NES ratios greater than 2-fold in all three replicates in at least three tissues for being nucleus-specific. All data for the identified proteins and ratios between samples is reported in Table S2.

